# Re-evaluating inheritance in genome evolution: widespread transfer of LINEs between species

**DOI:** 10.1101/106914

**Authors:** Atma M. Ivancevic, R. Daniel Kortschak, Terry Bertozzi, David L. Adelson

## Abstract

Transposable elements (TEs) are mobile DNA sequences, colloquially known as ‘jumping genes’ because of their ability to replicate to new genomic locations. Given a vector of transfer (e.g. tick or virus), TEs can jump further: between organisms or species in a process known as horizontal transfer (HT). Here we propose that LINE-1 (L1) and Bovine-B (BovB), the two most abundant TE families in mammals, were initially introduced as foreign DNA via ancient HT events. Using a 503-genome dataset, we identify multiple ancient L1 HT events in eukaryotes and provide evidence that L1s infiltrated the mammalian lineage after the monotreme-therian split. We also extend the BovB paradigm by increasing the number of estimated transfer events compared to previous studies, finding new potential blood-sucking parasite vectors and occurrences in new lineages (e.g. bats, frog). Given that these TEs make up nearly half of the genome sequence in today’s mammals, our results provide the first evidence that HT can have drastic and long-term effects on the new host genomes. This revolutionizes our perception of genome evolution to consider external factors, such as the natural introduction of foreign DNA. With the advancement of genome sequencing technologies and bioinformatics tools, we anticipate our study to be the first of many large-scale phylogenomic analyses exploring the role of HT in genome evolution.

**Significance statement:** LINE-1 (L1) elements occupy about half of most mammalian genomes (1), and they are believed to be strictly vertically inherited (2). Mutagenic L1 insertions are thought to account for approximately 1 of every 1000 random, disease-causing insertions in humans (4-7). Our research indicates that the very presence of L1s in humans, and other therian mammals, is due to an ancient transfer event – which has drastic implications for our perception of genome evolution. Using *a machina* analyses over 503 genomes, we trace the origins of L1 and BovB retrotransposons across the tree of life, and provide evidence of their long-term impact on eukaryotic evolution.

## Introduction

Transposable elements (TEs) are mobile segments of DNA which occupy large portions of eukaryotic genomes, including more than half of the human genome (1). Long interspersed element (LINE) retrotransposons are TEs which move from site to site using a “copy and paste” mechanism, facilitating their amplification throughout the genome (3, 4). The insertion of retrotransposons can interrupt existing genetic structures, resulting in gene disruptions, chromosomal breaks and rearrangements, and numerous diseases such as cancer (5-8). Two of the most abundant retrotransposon families in eukaryotes are LINE-1 (L1) and Bovine-B (BovB) (2, 9).

Horizontal transfer (HT) is the transmission of genetic material by means other than parent-to-offspring: a phenomenon primarily associated with prokaryotes. However, given a vector of transfer (e.g. virus, parasite), retrotransposons have the innate ability to jump between species as they do within genomes (3, 10). Studies investigating the possibility of HT in retrotransposons are limited, mainly including CR1s and RTEs (9, 11-13). Given the limited evidence to date, we tested the hypothesis that horizontal transfer is a ubiquitous process not restricted to certain species or retrotransposons. We used L1 and BovB elements as exemplars because of their contrasting dynamics and predominance in mammalian genomes. BovB retrotransposons provide an excellent example of horizontal transfer: divergent species contain highly similar BovB sequences and the analysis of various tick species reveals a plausible vector of transfer (9). In contrast, L1 elements are believed to be only vertically inherited, based on knowledge gained primarily through mammalian organisms (2). We hypothesise that the very presence of L1s in today’s mammals is due to an ancient HT event. In this study, we use BovBs as a comparison to identify common characteristics of horizontally transferred elements in contemporary eukaryotic species.

Three criteria are typically used to detect HT candidates: 1) a patchy distribution of the TE across the tree of life; 2) unusually high TE sequence similarity between divergent taxa; and 3) phylogenetic inconsistencies between TE tree topology and species relationships (14). To comprehensively test these criteria, we performed large-scale phylogenomic analyses of over 500 eukaryotic genomes (plants and animals) using iterative similarity searches of BovB and L1 sequences.

## Results

### Distribution and abundance of TEs across species

Our findings show that there are two phases in HT: effective insertion of the TE, followed by expansion throughout the genome. Figure 1 shows that both BovB and L1 elements have a patchy distribution across eukaryotes. Both are absent from most arthropod genomes yet appear in relatively primitive species such as sea urchins and sea squirts. Furthermore, both TEs are present in a diverse array of species including mammals, reptiles, fish and amphibians. The main difference between BovB and L1 lies in the number of colonised species. BovBs are only present in 60 of the 503 species analysed, so it is easy to trace their horizontal transfer between the distinct clades (e.g. squamates, ruminants). In contrast, L1s encompass a total of 407 species, within plants and animals, and are ubiquitous across the well-studied therian mammals. However they are surprisingly absent from several key species, including platypus and echidna (monotremes).

**Figure 1:**
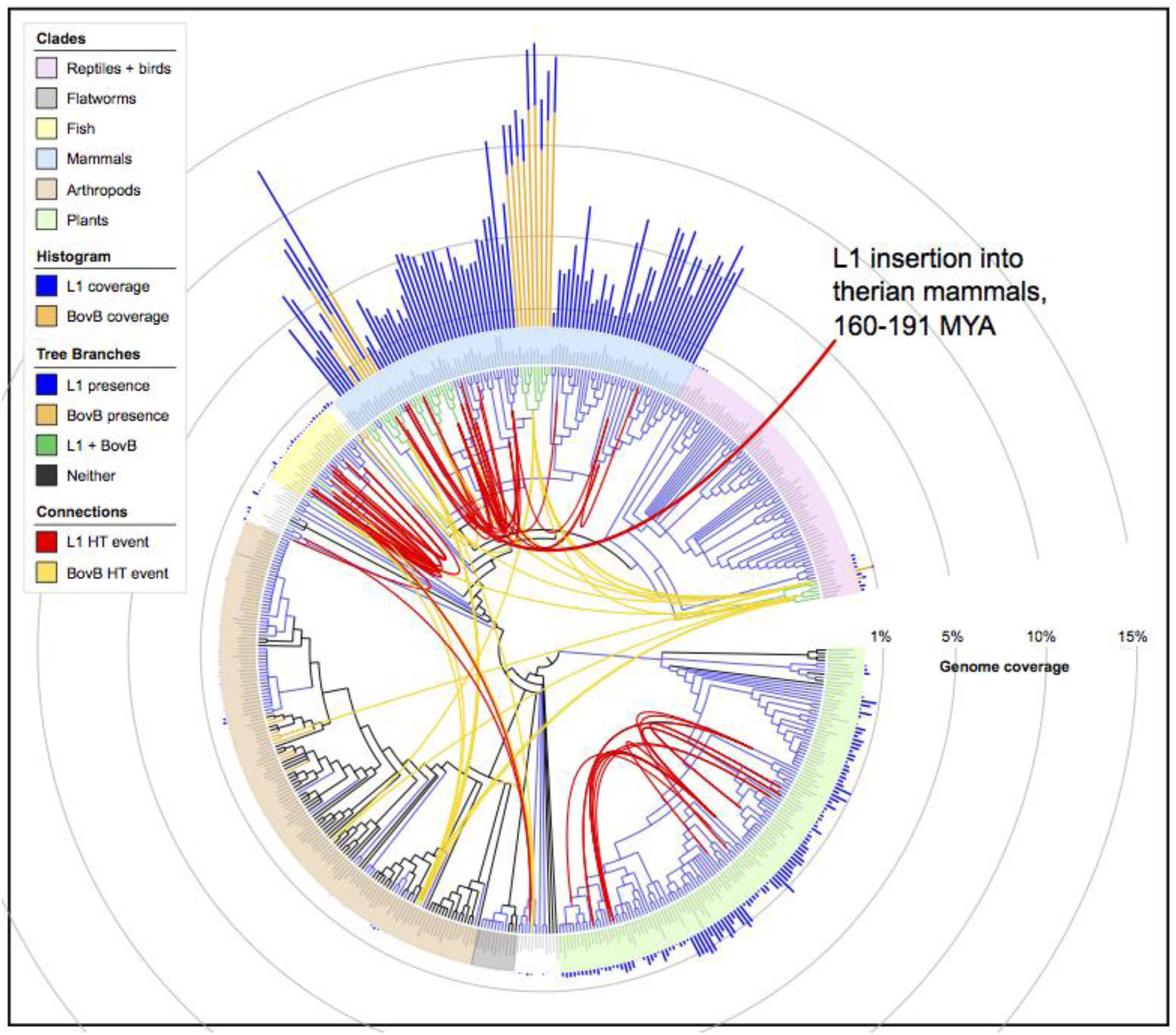
Presence and coverage of L1 and BovB elements across eukaryotes. The Tree of Life (51) was used to infer a tree of the 503 species used in this study; iTOL (52) was used to generate the bar graph and final graphic. The red arrow marks the L1 horizontal transfer event into therian mammals between 163-191 MYA. Branches are coloured to indicate which species have both BovB and L1 (green), only BovB (orange), only L1 (blue), or neither (black). Bar graph colours correspond to BovB (orange) and L1 (blue). Connections indicate possible HT events involving BovB (yellow) or L1 (red) elements. An interactive and downloadable version of this figure is available at: http://itol.embl.de/shared/atma

The abundance of TEs differs greatly between species. As shown in Fig. 1, mammalian genomes are incredibly susceptible to BovB and L1 expansion. More than 15% of the cow genome comprises these TEs (12% BovB, 3% L1). This is without considering the contribution of TE fragments (16) or derived Short INterspersed Elements (SINEs), boosting retrotransposon coverage to almost 50% in mammalian genomes (1). Even within mammals there are noticeable differences in copy number; for example, bats and equids have a very low number of full-length BovBs (<50 per genome) compared to the thousands found in ruminants and Afrotherian mammals. The low copy number here is TE-specific rather than species-specific; there are many L1s in bats and equids. Hence, the rate of TE propagation is determined both by the genome environment (e.g. mammal versus non-mammal) and the type of retrotransposon (e.g. BovB versus L1).

### BovB has undergone numerous, widespread HT events

To develop a method for identifying horizontal transfer events, we used BovB, a TE known to undergo HT. We clustered and aligned BovB sequences (both full-length nucleotide sequences and amino acid reverse-transcriptase domains) to generate a representative consensus for each species and infer a phylogeny (Fig 2a shows the nucleotide-based tree). The phylogeny supports previous results (9)—with the topology noticeably different from the tree of life (Fig. 1)—although we were able to refine our estimates for the times of insertion. For example, the cluster of equids includes the white rhinoceros, *Ceratotherium simum*, suggesting that BovBs were introduced into the most recent common ancestor before these species diverged. The low copy number in equids and rhinoceros, observed in Fig. 1, is not because of a recent insertion event; the most likely explanation being that the donor BovB inserted into an ancestral genome, was briefly active, but lost its ability to retrotranspose and was subsequently vertically transmitted.

**Figure 2:**
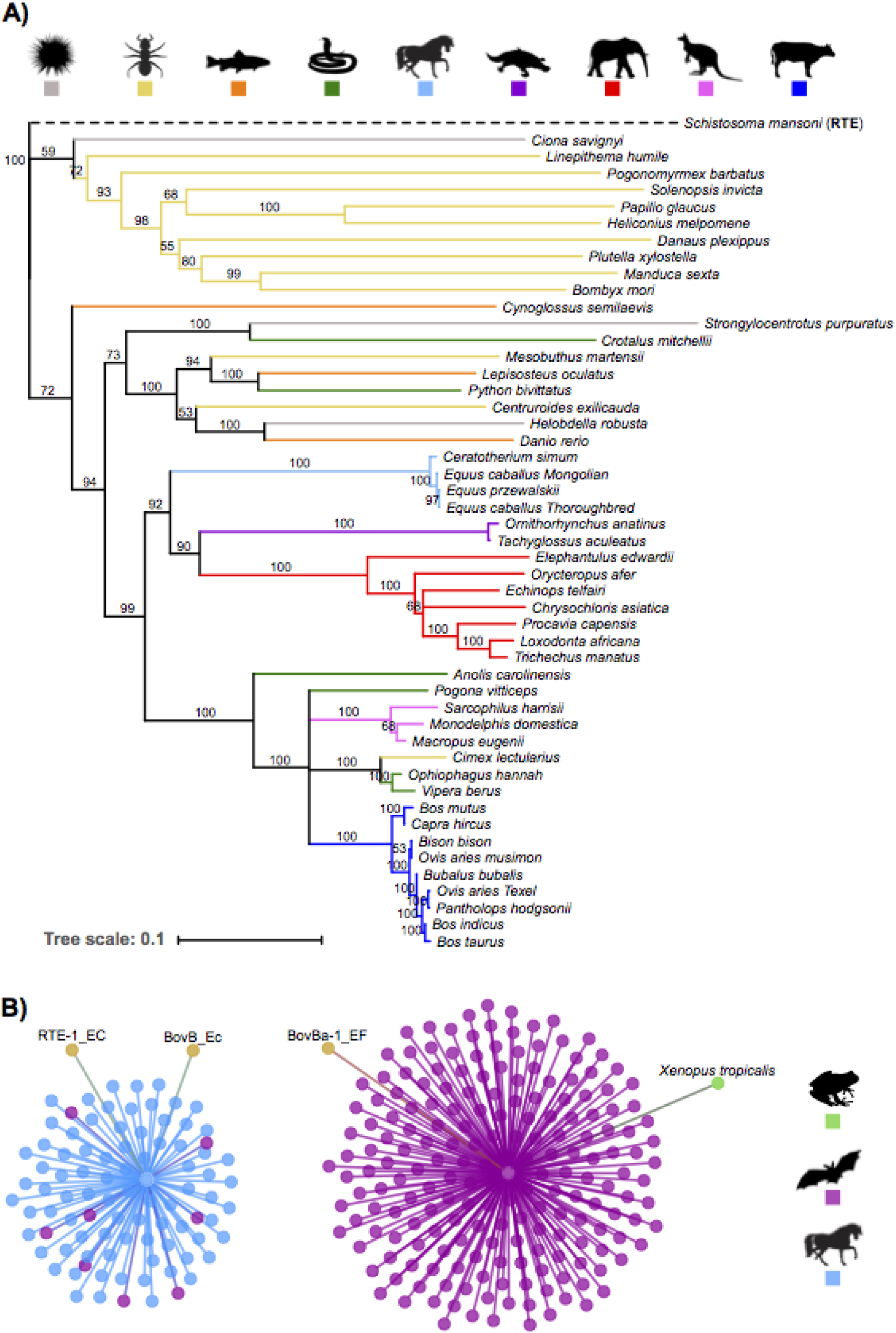
HT of BovB retrotransposons. (2a) Neighbour-joining tree (1000 bootstrap replicates) inferred using full-length nucleotide BovB consensus sequences, representing the dominant BovB family in each species. Nodes with confidence values over 50% are labelled and branches are coloured taxonomically. RTE sequence from *Schistosoma mansoni* was used as the outgroup. (2b) Network diagram representing the two distinct BovB clades in bats. Nodes are coloured taxonomically apart from the RepBase (17) sequences (light brown). RTE-1_EC and BovB_Ec are shown to belong to a single family, while BovBa-1_EF-like bat sequences form a separate family containing a single full-length BovB from the frog *Xenopus*.

The placement of arthropods is intriguing, revealing potential HT vectors and the origin of BovB retrotransposons. For example, BovBs from butterflies, moths and ants appear as a monophyletic group, sister to sea squirt *Ciona savignyi* BovB. The presence of BovB in all these species suggests that BovB TEs may have arisen as a subclass of ancient RTEs, countering the belief that they originated in squamates (13). The next grouping consists of two scorpion species (*Mesobuthus martensii* and *Centruroides exilicauda*) nestled within the snakes, fish, sea urchin and leech—a possible vector. But the most interesting arthropod species is *Cimex lectularius*, the common bed bug, known to feed on animal blood. The full-length BovB sequence from *Cimex* shares over 80% identity to viper and cobra BovBs; their reverse transcriptase domains share over 90% identity at the amino acid level. Together, the bed bug and leech support the idea (9) that blood-sucking parasites can transfer retrotransposons between the animals they feed on.

We extended the BovB analysis to include 10 bat species and one frog (*Xenopus tropicalis*). Bats were not included in the phylogenetic analysis because their BovB sequences were too divergent to construct an accurate consensus. Instead, we clustered all individual BovB sequences to identify two distinct subfamilies (Fig. 2b); one containing all the horse and rhino BovBs as well as eight bat sequences, and the other containing the remaining bat BovBs as well as the single BovB from *Xenopus*. We also included three annotated sequences from a public database (17) to resolve an apparent discrepancy between the naming of BovB/RTE elements. Our results have several implications: first, bat BovBs can be separated into two completely distinct clades suggesting bat BovBs arose from independent insertion events; second, the BovBa-1-EF bat clade may have arisen from an amphibian species, or vice versa; and third, the naming conventions used in RepBase (17) need updating to better distinguish BovB and RTE sequences. This third point is discussed in the Supplementary Information (see Supp. Fig. 1).

In order to exhaustively search for all cases of BovB HT, we tested several all-against-all clustering approaches to detect individual HT candidate sequences. We first replicated the method described in El Baidouri *et al*. (2014) (19), which uses BLAST (18) to compare all sequences within a database, and SiLiX (20) to extract discordant clusters. This worked well for recent BovB transfers (e.g. *Cimex lectularius*—snakes), but failed to identify ancient transfer events and required considerable computational time and power. Next, we tested VSEARCH (21), which is orders of magnitude faster than BLAST (18). 170,882 BovB sequences were clustered in under 14 minutes on a high computing cluster with 16 cores.

Many of the resulting clusters contained BovBs from closely related species (e.g. cow and yak). To find the most compelling HT events, we imposed the restriction that clusters had to contain BovBs from species that belonged to different eukaryotic Orders (e.g. Afrotheria and Monotremata). We performed *a machina* validation for each candidate HT cluster: pairwise alignments of the flanking regions to rule out possible contamination or orthologous regions, phylogenetic reconstructions to confirm discordant relationships, and reciprocal best hit checks to confirm correct clustering (see Methods). A total of 11 BovB clusters passed all of the tests; six cross-Phylum events and five cross-Class HT events (visualized in Figure 1; described in Supp. Table 6 and Supp. Figures 4-14).

**Figure 3:**
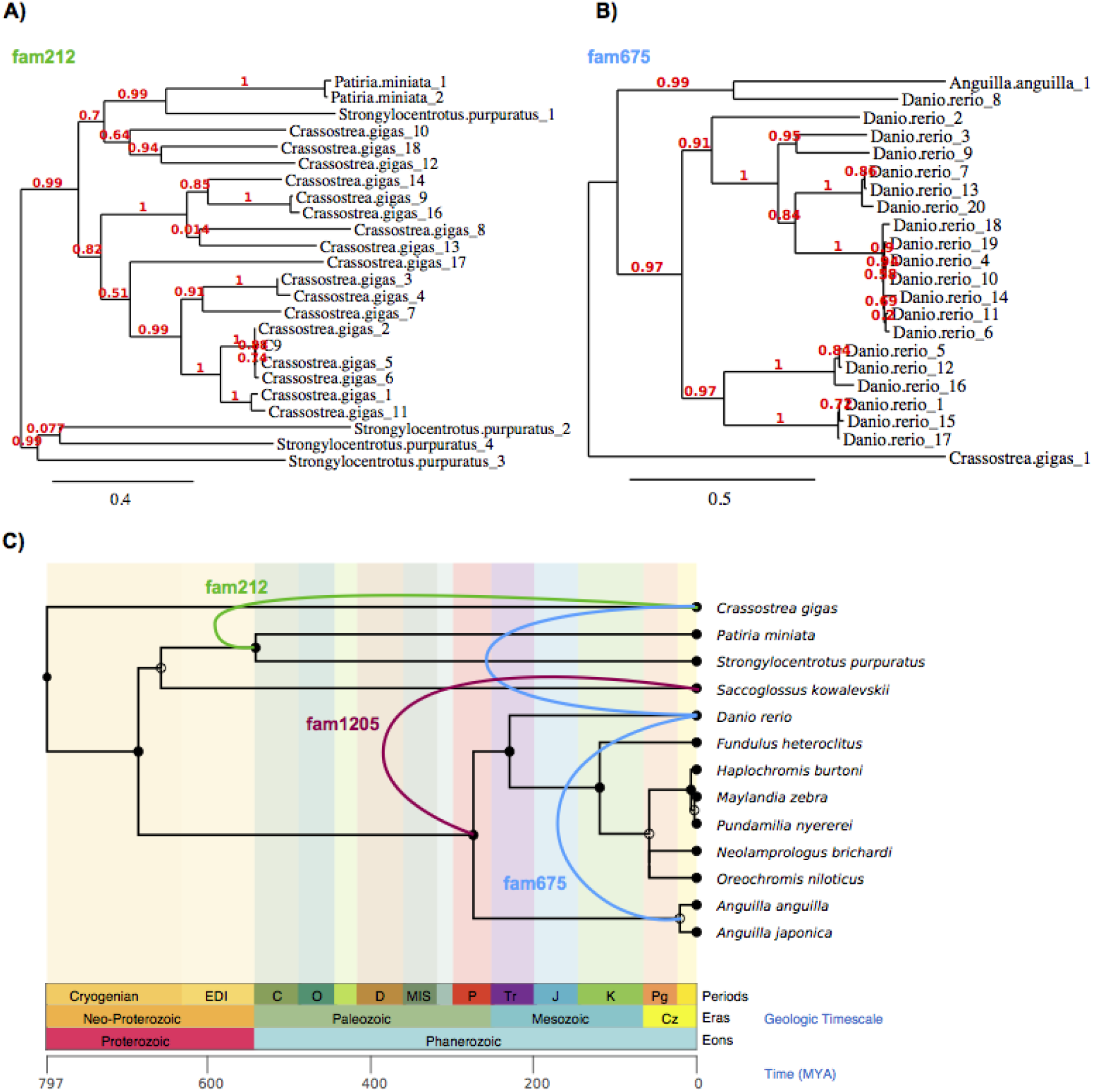
L1 cross-Phylum HT events. (3a) Phylogeny of putative cross-Phylum HT events involving ancient Tx1-like L1 elements in aquatic species. MUSCLE (46) was used to align the sequences, Gblocks (47) was used to extract conserved blocks from the alignment, PhyML (53) was used to infer a phylogeny and TreeDyn (54) was used for tree rendering. (3b) Same as (3a), for L1 fam675 involving zebrafish, eel and oyster. Additional discordant phylogenies are included in the Supplementary Information. (3c) TimeTree (55) illustrating the putative L1 horizontal transfer events. Shows only the species involved in the three cross-Phylum HTs. Background is coloured to match the ages in the geological timescale.

**Figure 4:**
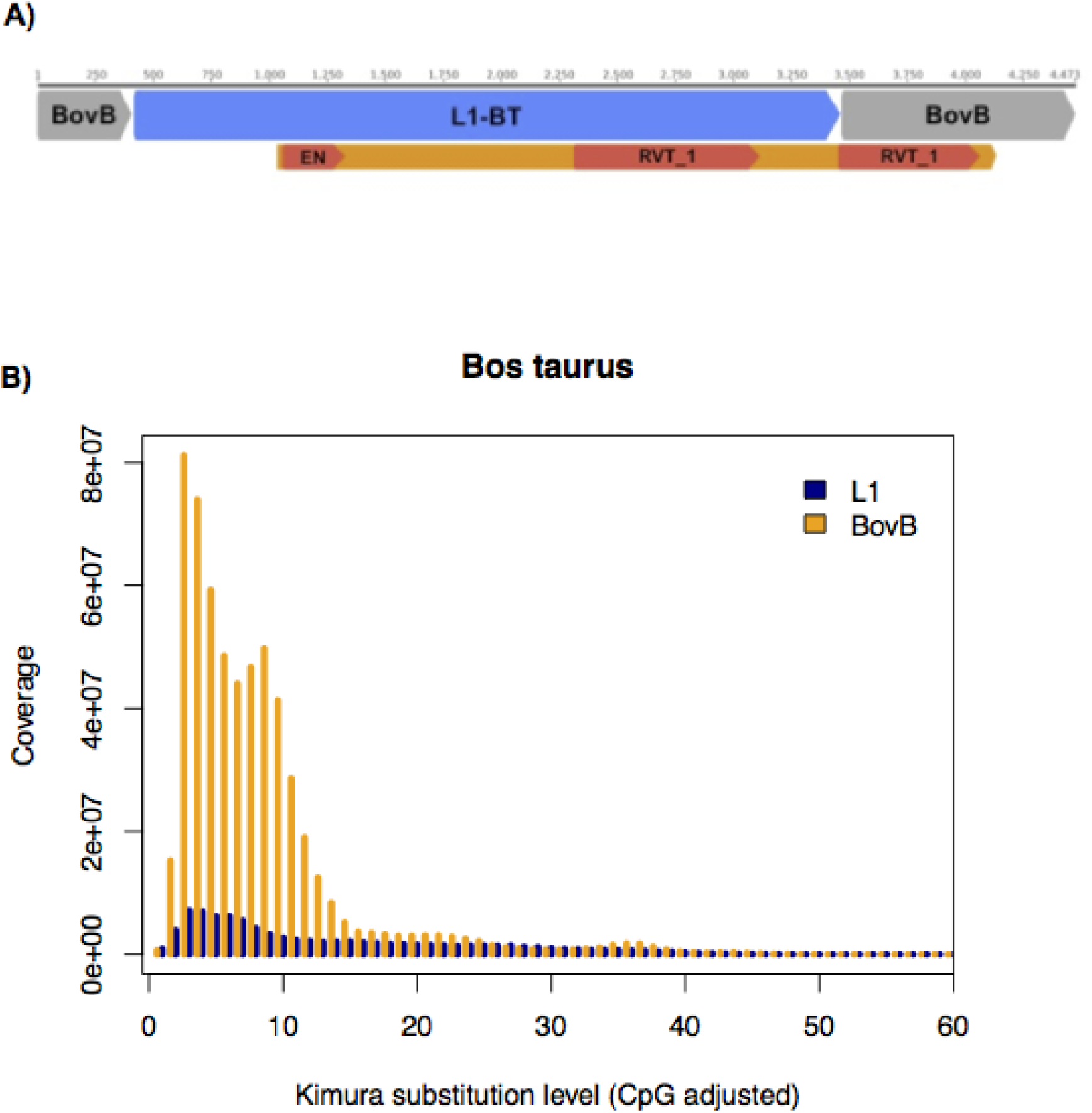
Chimeric L1-BovB element in cattle genomes. (4a) Chimeric L1-BovB retrotransposon found in cattle genomes (*Bos taurus* and *Bos indicus*). L1-BT and BovB correspond to RepBase names (17), representing repeats which are known to have been recently active. RVT_1 = reverse-transcriptase, EN = endonuclease domain. The orange bar is the length of the entire open reading frame. (4b) Kimura divergence plot of BovB and L1 elements in *Bos taurus*. RepeatMasker (50) was used to mask all BovB and L1 nucleotide sequences in the cow genome and calculate divergence from alignments to RepBase (17) consensus sequences. The y-axis represents coverage against the RepBase super consensus library; the x-axis indicates the Kimura divergence estimate. Ruminants are the only mammals which have currently active lineages of both BovB and L1 elements, explaining how the chimeric BovB-L1 element may have arisen.

The number of BovBs in each cluster was useful for inferring the direction of transfer. BovBs are thought to have entered ruminants after squamates (13). The clusters outlined in Supp. Table 6 show a recurring pattern where the ‘donor’ species has far fewer BovBs than the ‘acceptor’ species, indicating that BovB undergoes an expansion following the insertion event. For example, fam3 contains five BovBs from lizard *Pogona vitticeps* and 4008 BovBs from ruminant species. Likewise, fam4 contains 1 BovB from scorpion *Mesobuthus martensii* and 396 BovBs from Afrotherian species. This supports the theory that retrotransposons undergo HT to expand in new species, and escape suppression from their current host (19). Altogether, our results demonstrate that the horizontal transfer of BovB elements is even more widespread than previously reported, providing one of the most convincing examples of eukaryotic HT to date.

### L1 horizontal transfer is infrequent yet possible

We carried out the same exhaustive search in L1s, which presented a challenge because of greater divergence and the sheer number of vertically inherited copies. Producing a consensus for each species was impractical as most species contained a mixture of old degraded L1s and young intact L1s. Instead, we used the all-against-all clustering methods on the collated dataset of L1 nucleotide sequences over 3kb in length (>1 million sequences total). Once again, VSEARCH (21) was substantially faster and identified more potential HT candidates than the BLAST+SiLiX method (18-20). This is likely due to a crucial difference in clustering algorithms; SiLiX uses single linkage to draw connections between sequences, which is effective for recent HT events but clusters all “degraded” elements into a single group. In contrast, VSEARCH relies on centroid/average linkage, and is thus more appropriate for ancient HT events (where the centroid is the transferred TE).

Over 9000 clusters contained L1s from at least two different species: these were our HT candidates. As before, to improve recognition of HT, we looked for families displaying cross-Order, cross-Class or cross-Phylum transfer. We checked for elevated sequence identity compared to flanking regions; more than two L1 sequences in each cluster; recognizable open reading frames or reverse-transcriptase domains; discordance in phylogenetic reconstructions or absence in neighboring species; and the presence of fragments in the genome to support each HT candidate. By using the procedure we had established for BovB elements, we were able to retain 39 L1 clusters as potential HT events (illustrated in Figure 1; described in Supp. Table 7 and Supp. Figures 15-31).

Three clusters portray ancient cross-Phylum L1 HT events. Two of these (Figure 3a-c) include oyster (*Crassostrea gigas*) as the potential vector species. The third cluster (Supp. Figure 15, Figure 3c) involves various fish species and *Saccoglossus kowalevskii*, a type of worm typically found in marine sediments. The L1 sequences in all three cross-Phylum clusters have intact reverse transcriptase domains and resemble the diverse Tx1-like L1s originally discovered in *Xenopus* frogs (15, 22). The cross-Class cluster (fam35 in Supp. Table 7, Supp. Figure 16) suggests a transfer event between ray-finned fishes and coelacanth (*Latimeria chalumnae*). A further nine clusters depict cross-Order L1 HTs between different fish species; many of which include zebrafish (*Danio rerio*) and killifish (*Fundulus heteroclitus)*. Taken together, these results suggest that L1s prefer aquatic species as intermediary species vectors for HT. It may also explain why the zebrafish genome (*Danio rerio*) has such a diverse repertoire of TEs, with multiple co-existing active L1 lineages (Supp. Figure 78) (23).

The remaining cross-Order L1 events involve plant and mammal groupings (Supp. Table 7). Among plants, frequently implicated Orders include Solanales (flowering plants such as tomato and potato), Fagales (trees *Betula nana, Castanea mollissima*), and Vitales (grapevine *Vitis vinifera*). Like zebrafish, *Vitis vinifera* contains an unusual array of L1 lineages (see Supp. Figures 18, 21); some of which are potentially still active based on their structural integrity (15).

The mammalian cross-Order L1 events were hardest to reconcile due to sequence degradation and an overwhelming amount of noise from vertically inherited copies. Many clusters include representatives from Perissodactyls (horses, rhinoceros), Cetartiodactyls (hoofed animals and whales) and Chiroptera (bats). The number of elements in a cluster mimicked the patterns seen with BovBs: very few elements from the ‘donor’ species, and an expansion of L1s in the ‘acceptor’ species (e.g. Supp. Table 7: fam97, fam165, fam5210, fam3167). However, many of the clusters simply contained a sporadic assortment of different mammals - potentially clustered together because they all contain “dead", inactive L1s. At this stage, we cannot confirm whether the mammalian clusters are due to ancient HT events or similarities based on accumulated mutations and extinct L1 subfamilies.

Finally, our mining of L1 HT candidates led to the serendipitous discovery of a chimeric L1-BovB element present in cattle genomes (*Bos taurus* and *Bos indicus*), shown in Fig. 4a. This particular element most likely arose from a recently active L1 element (98% identical to the canonical *Bos* L1-BT (17)) inserting into an active BovB (97% identical to *Bos* BovB (17)). Ruminants are the only mammals that currently have active lineages of both BovB and L1 elements (Figure 4b). Such a vast and ongoing expansion of TEs has created the ideal genomic environment for the genesis of chimeric repetitive elements. With two reverse-transcriptase domains and high similarity to active L1/BovB elements, this particular chimeric element could still be functional. This raises the question - can L1 elements to be transferred throughout mammals by being transduced by BovBs?

## Discussion

### The curious case of L1 absence in monotremes

Figure 1 shows the similarly patchy distributions of BovB and L1 elements across our inferred tree of life. Monotremes are a particularly interesting discrepancy because they contain BovBs, yet lack L1s. There are several possible explanations for this: either L1s could not be detected due to the draft status of the genomic data; or L1s were expunged shortly after the monotreme-therian split but before they had a chance to accumulate; or monotremes never had L1s. To control for genome quality, we used two independent searching strategies to mine for L1s in both full genome data and all available nucleotide databases, as well as a third method to act as a reciprocal best-hit check (see Material and Methods). Species were annotated ‘L1-present’ if there was any evidence of fragments or full-length copies from at least one of the methods. There was no hit at all for echidna, and the few isolated fragments found in the platypus assembly are known contaminants from wallaby (15). We could easily identify other TE families in both species, including an abundance of L2s and BovBs.

The second scenario is also unlikely in the context of L1 distributions in other eukaryotes. TE removal from a genome is thought to occur through a series of large (>10kb) segmental duplications (24). However, this process is not absolute; it is difficult to remove all evidence of a TE family, especially since extinct and degraded copies are unlikely to carry a selective disadvantage. Consider the 60 analysed bird genomes: full-length L1s have been eradicated from the avian lineage, yet every bird species bears evidence of ancient/ancestral L1 activity through the presence of fragments (15). Similarly, L1s have been functionally inactive in megabats for at least 24 million years (25), yet their genomic history is preserved via degraded L1 remains. This is not the case for platypus or echidna. We therefore conclude that L1s were never present in the monotreme lineage. To emphasize this, the L1 explosion in therian mammals mimics the rapid BovB expansions in ruminants and Afrotherian mammals (Figure 1). Our results indicate that L1s were inserted into a common ancestor of therian mammals 160 and 191 Million Years Ago (MYA), and have since been vertically inherited, with possible ongoing inter-mammal HTs.

### Both BovB and L1 satisfy the criteria for horizontal transfer

The typical criteria used to infer horizontal transfer are a patchy distribution across taxa, phylogenetic inconsistencies in the TE topology, and high TE sequence similarity between divergent species. As discussed above, both BovB and L1 have a patchy distribution across the eukaryotic tree of life. Both BovB and L1 also show phylogenetic inconsistencies in TE topology – for BovB, this is evident from just the consensus tree (Figure 2a); for L1, this is shown in the individual phylogenies constructed for the HT clusters (Figure 3; Supp. Figures 15-31). In each case, the species involved in the HT event appear too closely related, and neighbouring species lack evidence of similar copies.

In terms of high sequence similarity, the level of identity between transferred elements seems largely dependent on how recently the HT event occurred. For example, consider the BovB HT events. The BovB element from bed bug *Cimex lectularis* shares over 80% similarity to BovBs from three snake species (Figure 2a, Supp. Figure 4), suggestive of a recent event. Ancient HT events are unlikely to have such a high degree of similarity, due to accumulated mutations over time. In BovB, the ancient HT events were found by reducing the clustering identity to 50-60% and using a centroid-based clustering strategy (21) rather than single linkage (20).

The L1 HT families satisfy these same criteria. Using stringent identity parameters of 80% or higher, we could not find any candidates; there have been no recent L1 HT events in our subset of species. However, the ancient L1 transfers between aquatic species, plants, and even mammals (Figure 3, Supp. Table 7) mimic the ancient BovB events. The sequence similarity is restricted to the TE sequence, and the ‘donor’ species usually has fewer elements than the new host species. This contradicts the belief that L1s are exclusively vertically inherited, and supports our conclusion that a similar event introduced L1s to mammals.

### Transfer frequency and mechanisms differ between TE classes

The main argument against L1 HT is the frequency of transfer versus number of colonised species. For example, consider the number of cross-Phylum HT events found for each TE. BovBs have undergone at least six cross-Phylum transfers (between widely divergent groups such as reptiles and mammals), yet they are only present in 60 of the 503 analysed species. In contrast, we were only able to find evidence for three cross-Phylum L1 HTs (all among sea dwelling creatures). These three events cannot explain how L1s arose in therian mammals, or came to colonise over 400 species. If L1 HT is so rare, how have L1s come to dominate almost all of the major clades of plants and animals?

There are several explanations for these observations. First, L1s are ancient: they have been around for millions of years longer than BovBs, which only emerged recently (possibly as a subclass of ancient RTEs). BovB HT is easy to trace because we can see the likely insertion point for each distinct group of species (Figure 1). In contrast, L1 HT events potentially occurred before the origin of today’s species. If L1s inserted into an early ancestor of therian mammals, it is also possible they inserted into the ancestor of sauropsids, or fish. The patterns we observe in Figure 1 could then be explained by subsequent vertical inheritance into descendent species.

Another aspect to consider is the mechanism of transfer. BovB and L1 are similar in structure, but L1 sequences are almost twice as long as BovB. This may reduce the likelihood of successful transfer and integration into the new host genome. Moreover, it seems unlikely that L1s are carried via parasitic insects, like BovBs. The cross-Phylum clusters implicate basal metazoans such as oysters, molluscs and marine worms as possible vectors of L1 HT. Compared to arthropods, these types of species are heavily underrepresented in our dataset; we are missing numerous potential intermediary species. Further studies should explore different vectors of transfer (microbiological or viral) to provide a more comprehensive representation of the tree of life.

Finally, our analysis only considered TE candidates from cross-Order species, to find the most extreme cases of HT. Several studies have suggested that HT is more likely to occur between closely related species with similar genomic environments (11, 26, 27). The BovB results (e.g. Figure 2a, Supp. Figures 4-14) suggest that there has been ongoing HT even between ruminant species. Identifying HT events between similar species or individuals of the same species would give a better approximation for TE transfer frequency.

### Suppression of L1 and BovB activity in megabat genomes

Both the BovB and L1 results suggest that transferred TEs can retain activity and expand within their new host. However, the extent to which a TE can propagate in a new organism depends on factors such as a favourable genomic environment and TE replication machinery. Mammals are generally more susceptible to TE expansion than other species; this is evident in Figure 1. However, bats appear to be exceptional in their ability to quickly suppress LINE activity.

Bats, particularly megabats, are often used as an example of L1 extinction affecting an entire lineage (15, 25). Bat BovB sequences are similarly degraded (Supp. Figures 43-53). Despite the presence of two BovB subfamilies, indicative of two independent HT events, bat BovBs show little evidence of replication and no preserved functional domains. In fact, the megabat group is the only monophyletic clade on our tree showing L1/BovB presence coupled with complete extinction of both TE families. This is important in the context of host suppression mechanisms.

Bats are frequently implicated as vectors of DNA exchange: they transmit numerous viruses and TEs, cause disease epidemics, and feed on arthropods (28-30). As such, they are the ideal intermediate species for horizontal transfer. Constant exposure to potentially harmful DNA may have led to the evolution of heightened TE silencing mechanisms. This is supported by the observation that bats have a relatively compact genome size, and experience dynamic loss and gain of DNA (24). It is likely that bats act as TE reservoirs (31): enabling the transmission of foreign DNA while minimizing impact to their own genomes.

### HT of L1s potentially catalyzed the evolution of therian mammals

Over 30 years ago, Barbara McClintock pioneered the discovery of transposable elements, flagging them as “controlling elements” of the genome (32). In the last few years, we have finally started seeing evidence of their functional importance. A study of 29 mammals found over 280,000 non-coding elements exapted from TE insertions (33). TEs have been implicated in the evolution of innate immunity (34, 35) and the placenta (36, 37), as well as transcriptional regulation of mammalian brains (38). The structural changes arising from horizontally transferred TEs have contributed to the modification of regulatory elements and led to the development of novel traits (recently reviewed by Boto (39)).

Perhaps most intriguingly, recent evidence shows that Krüppel-associated box domain-containing zinc-finger proteins (KRAB-ZFPs) use TEs, particularly endogenous retroviruses and L1s, to establish species-specific networks of epigenetic regulation (40, 41). Humans have essentially domesticated L1 elements to the extent that transposon-based regulatory sequences are fundamental for various developmental and physiological pathways. If L1s were transferred into therian mammals as foreign DNA, the consequent L1 expansion, domestication and modification of regulatory networks may explain the rapid speciation that occurred following the split from monotremes. Importantly, it demonstrates that HT of TEs can have drastic and lasting effects on new host genomes, emphasizing the stochasticity of genome evolution.

## Conclusions and implications

Our analyses indicate that both BovB and L1 retrotransposons can undergo HT, albeit at different rates. We extracted millions of retrotransposon sequences from a 503-genome dataset, demonstrating the similarly patchy distributions of these two LINE classes across the eukaryotic tree of life. We further extended the analysis of BovBs to include blood-sucking arthropods capable of parasitizing mammals and squamates, as well as two distinct clades of bat BovBs and the first report of BovB in an amphibian. Contrary to the belief of exclusive vertical inheritance, our results with L1s suggest multiple ancient HT events in eukaryotes, particularly among aquatic species, and HT into the early therian mammal lineage. The rapid speciation following the split of theria and australosphenids (monotremes), between 160-191 MYA, coincides with the invasion of L1 elements into therian genomes. We therefore speculate that the speciation of therian mammals was driven in part by the effect of L1 retrotransposition on genome structure and function (including regulatory effects on transcriptional networks). This ancient transfer event allowed expansion of L1s and associated SINEs, transformation of genome structure and regulation in mammals (8) and potentially catalysed the therian radiation.

## Materials and Methods

Detailed description of the methods, including tables and figures, are available in the Supplementary Information. Sample code for each step is available at: https://github.com/AdelaideBioinfo

### Extraction of L1 and BovB retrotransposons from genome data

To extract the retrotransposons of interest, we used the methods and genomes previously described in Ivancevic *et al.* (2016) (15). Briefly, this involved downloading 499 publicly available genomes (and acquiring four more from collaborations), then using two independent searching strategies (LASTZ (42) and TBLASTN (18)) to identify and characterise L1 and BovB elements. A third program, CENSOR (43), was used with the RepBase library of known repeats (17) to verify hits with a reciprocal best-hit check. The raw L1 results have been previously published in Ivancevic *et al*. (2016) (15) (Supplementary Material); the BovB results are included in the Supplementary Information.

### Extraction and clustering of conserved amino acid residues

Starting with BovBs, USEARCH (44) was used to find open readings frames (ORFs), with function -fastx_findorfs and parameters -aaout (for amino acid output) and -orfstyle 7 (to allow non-standard start codons). HMMer (45)was used to identify reverse transcriptase (RT) domains within the ORFs. RT domains were extracted using the envelope coordinates from the HMMer domain hits table (-domtblout), with minimum length 200 amino acid residues. The BovB RT domains from all species were collated into one file and clustered with UCLUST (44). This was done as an initial screening to detect potential horizontal transfer candidates. The process was repeated with L1 elements.

### Clustering of nucleotide sequences to build one consensus per species

The canonical BovB retrotransposon is 3.2 kb in length (9, 17), although this varies slightly between species. In this study, we classified BovB nucleotide sequences ≥2.4kb and ≤4kb as full-length. We wanted to construct a BovB representative for each species. Accordingly, for each species, UCLUST (44) was used to cluster full-length BovB sequences at varying identities between 65-95%. A consensus sequence of each cluster was generated using the UCLUST -consout option.

The ideal cluster identity was chosen based on the number and divergence of sequences in a cluster. E.g. for species with few BovBs, a lower identity was allowed; whereas for species with thousands of BovBs, a higher identity was needed to produce an alignable cluster. The final clustering identity and cluster size for each species is given in Supp. Table 1. Note that the bat species are not included in this table - they were clustered separately, due to the high level of divergence between BovBs.

This method was tested on L1 retrotransposons, but the results were not ideal; most species simply had too many L1 sequences. Other methods tested on both BovBs and L1s included using centroids instead of consensus sequences (this gave better alignments but was less representative of the cluster), and using the same clustering identity for all species (e.g. 80% - this did not work well for species with less than 100 elements in the genome).

### Inferring a phylogeny from consensus sequences

Consensus sequences were aligned with MUSCLE (46). The multiple alignment was processed with Gblocks (47) to extract conserved blocks, with default parameters except min block size: 5, allowed gaps: all. FastTree (48) was used to infer a maximum likelihood phylogeny using a general time reversible (GTR) model and gamma approximation on substitution rates. Geneious Tree Builder (49) was used to infer a second tree using the neighbour-joining method with 1000 bootstrap replicates.

### Distinguishing RTEs from BovBs

All sequences which identified as BovB or RTE were kept and labelled accordingly to their closest RepBase classification (17). However, there appeared to be numerous discrepancies with the naming: e.g. some RTE sequences shared >90% identity to BovBs, and vice versa.

BovB retrotransposons are a subclass of RTE, and they were only discovered relatively recently. It is likely that several so-called RTE sequences are actually BovBs.

To determine which species had BovB sequences, and which only had RTEs, we used the species consensus approach to build a BovB/RTE phylogeny (see Supp. Figure 1). This effectively separated BovB-containing species from RTE-containing species. The RTE sequences were not included in further analyses.

### Clustering of nucleotide BovB sequences from bats and *Xenopus*

A reliable BovB consensus could not be generated for any of the ten bat species because the sequences were too divergent and degraded. Some bat BovBs seemed similar to equid BovBs; others did not. Likewise, the single full-length BovB from frog *Xenopus tropicalis* was very different to canonical BovBs, sharing highest identity with the bats.

In an effort to characterise these BovBs into families, we grouped all full-length BovB sequences from the bats, frog, equids and white rhino into a single file. We also added two RepBase equid sequences (RTE-1_EC and BovB_Ec) and 1 RepBase bat sequence (BovBa-1_EF) (17). After clustering, we expected to find one family of equid BovBs, the equid RTE sequence as an outlier, and numerous families containing bat and frog BovBs.

The actual findings are described in the text (Fig. 2b). We used UCLUST (44) to cluster the sequences (function -cluster_fast with parameters -id, -uc, -clusters). The highest identity at which there were only 2 clusters/families was 40%. At higher identities, the equid BovBs stayed together but the bat and frog BovBs were lost as singletons.

To confirm the clustering, we also used MUSCLE to align all the sequences and FastTree to infer a phylogeny (see Supp. Figure 2).

### HT candidate identification - BovB and L1

We compiled all confirmed BovB and L1 sequences into separate multi-fasta databases (170,882 and 1,048,478 sequences respectively). The length cut-off for BovBs was ≤2.4kb and ≥4kb; for L1s, ≥3kb. BovBs were analysed first to identify characteristics of horizontal transfer events.

To detect HT candidates, we initially used the all-against-all clustering strategy described in El Baidouri*et al*. (2014) (19). Briefly, this method used a nucleotide BLAST (18) to compare every individual sequence in a database against every other sequence; hence the term all-against-all. BLAST parameters were as follows: -r 2, -e 1e-10, -F F, -m 8 (for tabular output). The SiLiX program (20) was then used to filter the BLAST output and produce clusters or families that met the designated identity threshold.

This method worked well for recent HT events (e.g. transfer between bed bug and snakes), but failed to pick up ancient HT events. For comparison, we also tested VSEARCH (21): an open source version of USEARCH that is orders of magnitude faster than BLAST (18) and uses centroid/average linkage to identify clusters. As before, we used our entire database of BovB sequences as input to VSEARCH, at clustering identities of 40-90%.

The majority of clusters contained several copies of the same BovB family from a single species - indicative of vertical inheritance. We found that using a lower identity threshold was more informative for capturing ancient HT events. At 50% identity, the clustering preserved the recent, high-identity HT events while also finding the ancient, lower-identity HT events. We concluded that this was the best % identity to use for our particular dataset, considering it includes widely divergent branches of Eukaryota.

Clusters were deemed HT candidates if they contained BovB elements belonging to at least two different species. To reduce the number of possible HT clusters, we went one step further and kept only the clusters which demonstrated cross-Order transfer (e.g. BovBs from Monotremata and Afrotheria in the same cluster). All potential HT candidates were validated by checking that they were not located on short, unplaced scaffolds or contigs in the genome. The flanking regions of each HT candidate pair were extracted and checked (via pairwise alignment) to ensure that high sequence identity was restricted to the BovB region. This was done to check for contamination or orthologous regions. Phylogenies of HT candidate clusters were inferred using maximum likelihood and neighbour-joining methods (1000 bootstraps).

The same procedure was performed to screen for nucleotide L1 HT candidates. VSEARCH (21) again proved to be more appropriate for finding HT events. As an extra step for L1s, we also used all ORF1 and ORF2 amino acid sequences from a previous analysis (15) to conduct similar all-against-all searches. However, the amino acid clusterings did not produce any possible HT candidates.

### Kimura divergence estimates for species containing both TEs

From our 503-genome dataset, we identified 47 species containing both BovB and L1 elements. To compare TE dynamics within these species, we used RepeatMasker (50) to compare L1 and BovB nucleotide sequences from each genome against the super consensus library of repeats curated by RepBase (17). Kimura substitution levels were calculated from the alignments using the provided RepeatMasker utility scripts (50). Supp. Figures 32-83 show the resulting plot for 46 species (*Bos taurus* plot appears as Figure 4b in the main text).

## Acknowledgements

The authors wish to acknowledge Olivier Panaud and Steve Turner for helpful discussions, Reuben Buckley and Lu Zeng for ideas and moral support, Brittany Howell for proof reading and providing a much-needed sanity check and Matt Westlake for HPC support above and beyond the call of duty.

## Author contributions

A.M.I. performed the analysis, interpreted the results and wrote the manuscript. R.D.K, T.B. and D.L.A. supervised the development of work and assisted in analysing the results and writing the manuscript. T.B. provided access to DNA samples and performed laboratory validation experiments.

The authors declare no conflict of interest.

## Supplementary information

Additional supplementary information is provided in the attached PDF.

## Data availability

Data generated or analysed during this study are included in the main text and Supplementary Information. Raw sequences are provided upon request.

